# SARS-CoV-2 Infects Peripheral Sensory Neurons and Promotes Axonal Degeneration via TRPV1 Activation

**DOI:** 10.1101/2025.03.06.641885

**Authors:** Vanessa Anna Co, Siwen Liu, Rachel Chun-Yee Tam, Bobo Wing-Yee Mok, Honglin Chen, Yiling Hong

**Author notes:** **Corresponding authors: Honglin Chen**, Professor, State Key Laboratory for Emerging Infectious Diseases & Department of Microbiology, Li Ka Shing Faculty of Medicine, and Centre for Virology, Vaccinology and Therapeutics, The University of Hong Kong, 21 Sassoon Road, Pokfulam, Hong Kong SAR, PR China. **Yiling Hong**, Professor, College of Veterinary Medicine, Western University of Health Sciences, 309 East Second Street, Pomona, CA 91766-1854, USA.

## Abstract

Common neurological symptoms of COVID-19, such as anosmia, headaches, and cognitive dysfunction, depend on interactions between the peripheral and central nervous systems. However, the molecular mechanisms by which SARS-CoV-2 affects the peripheral nervous system remain poorly understood, with ongoing debate about whether sensory neurons can be directly infected by the virus. In this study, human iPSC-derived sensory neurons were exposed to the SARS-CoV-2 BA.5 variant, a mutant virus, or viral S1 proteins. Under control conditions, sensory neurons exhibited low expression of ACE2. However, exposure to BA.5 or S1 proteins significantly upregulated ACE2 expression in peripherin-positive sensory neurons. Virological analysis confirmed that SARS-CoV-2 directly infects TRPV1-expressing sensory neurons, including olfactory neurons. Moreover, exposure to the live virus or S1 proteins induced TRPV1 upregulation and translocation from the nucleus to the cytosol, resulting in axonal destruction. Single-nucleus transcriptomic analysis revealed that viral exposure enhanced cAMP signaling, virus receptor and transmembrane transporter activities, and inflammatory regulation of TRP channels, which collectively contributed to synaptic and axonal damage. Importantly, treatment with a TRPV1 antagonist demonstrated neuroprotective effects. These findings underscore the need for further research into the interaction between SARS-CoV-2 and TRPV1, as well as its downstream signaling pathways, to develop therapeutic strategies for preventing sensory neuron loss during viral infections.

**Highlights:**

- iPSC technology was employed to generate peripheral sensory neurons from human induced pluripotent stem cells (iPSCs), providing a valuable platform for studying the impact of SARS-CoV-2 on peripheral sensory neurons.
- Our findings demonstrated that the SARS-CoV-2 Omicron BA.5 variant exerted both direct and indirect effects on peripheral sensory neurons. Virus exposure upregulated the angiotensin-converting enzyme 2 (ACE2) receptor in peripherin-positive neurons. Additionally, exposure to the virus or its S1 spike protein increased transient receptor potential vanilloid 1 (TRPV1) expression and trafficking, leading to axonal degeneration.
- Single-nucleus RNA sequencing revealed that BA.5 exposure enhanced cAMP signaling pathway, virus receptor and transmembrane transporter activities, and inflammatory regulation of TRP channels that led to the significantly damage on synapses and axon guidance.
- The TRPV1 antagonist capsazepine inhibited TRPV1 activation and mitigated axonal damage, offering neuroprotective effects for sensory neurons exposed to SARS-CoV-2.

## Introduction

The emergence of severe acute respiratory syndrome coronavirus 2 (SARS-CoV-2) triggered the coronavirus disease 2019 (COVID-19) pandemic ^1^. In addition to impacting the pulmonary system, infection with SARS-CoV-2 is commonly associated with neurological symptoms such as anosmia, headaches, and cognitive dysfunction ^2, 3^. Approximately 59% of COVID-19 patients developed signs of neuropathy in the mid-or long-term ^4^. Recent autopsy studies of patients who died with COVID-19 have indicated that SARS-CoV-2 is widely distributed within multiple respiratory and non-respiratory tissues, including the brain ^5^. Additionally, the virus has been detected in the olfactory cortex in a non-human primate model ^6^. While many respiratory viruses exhibit neurotropic properties, SARS-CoV-2 is notably association with neural impairment, particularly olfactory disorders ^7^. However, the molecular mechanisms underlying the impact of SARS-CoV-2 on the peripheral nervous system, the network of nerves that enables communication between the brain and the rest of the body, are not well understood. There are discrepancies in studies regarding the susceptibility of sensory neurons to SARS-CoV-2 infection. Some reports suggest that neurons may not be very susceptible due to their low expression of angiotensin-converting enzyme 2 (ACE2), the primary receptor for SARS-CoV-2 virus attachment and fusion, implying that neuropathies may result from indirect mechanisms affecting sensory neurons ^8, 9^. In contrast, other research indicates that sensory neurons express sufficient ACE2 and can indeed be infected by SARS-CoV-2 ^10^.

Transient receptor potential vanilloid 1 (TRPV1) is a calcium-permeable ion channel predominantly expressed in primary sensory neurons with unmyelinated C-fibers located the dorsal root ganglia (DRG) and trigeminal ganglia (TG) ^11^. TRPV1 plays a crucial role in several physiological processes, including intracellular calcium homeostasis, cellular chemotaxis, axonal guidance and neurite extension, cell proliferation and differentiation, and immune response ^12-15^. Additional, TRPV1 functions as an “attack sensor” under adverse conditions ^16^. TRPV1 activation involves phosphorylation by protein kinases and subsequent trafficking to the cytosol and membrane, where it becomes fully functional ^16-18^. Upregulation of TRPV1 expression on the plasma membrane enhances nociceptive responses and promotes release of pro-inflammatory neuropeptides that initiate a cascade of neurogenic inflammation ^19^. Furthermore, studies monitoring TRPV1 aggregation status, membrane mobility, and interaction with microtubules indicate that TRPV1 activation can rapidly disassemble dynamic microtubules, significantly affecting axonal growth, morphology, and migration ^20^. TRPV1 activation increases Ca^2^+influx, triggering plasma membrane depolarization and leading to cell death ^21-23^. Dysregulation of TRPV1 has been linked to the development of various neurological diseases ^24^. TRPV1 expression is reported to be significantly upregulated by several viral infections, including respiratory viruses such as human respiratory rhinovirus (HRV) and respiratory syncytial virus (RSV), as well as non-respiratory viruses like measles virus (MV) and hepatitis C virus (HCV) ^25, 26^. However, the interaction between SARS-CoV-2 and TRPV1 receptors remains largely unexplored.

To better understand the interactions between SARS-CoV-2 and sensory neurons, we examined SARS-CoV-2 infection in human sensory neurons derived from human iPSCs. Sensory neurons were infected with live viruses or exposed to the S1 subunit of the viral spike protein. The S1 protein facilitates viral attachment and fusion with the host cell membrane ^27^. SARS-CoV-2 S1 spike protein exposure has been shown to activate airway sensory C-fibers ^28^. We observed that approximately 5-10% of sensory neurons in our culture were directly infected by the virus. Additionally, following exposure to the virus and S1 protein, most TRPV1-positive sensory neurons showed upregulation and activation of the TRPV1 receptor.

## Results

### 1. Exposure to Omicron BA.5 Virus and Its Spike Protein S1 Subunit Upregulates ACE2 Expression in iPSC-Derived Peripherin-Positive Sensory Neurons

Peripheral sensory neurons were differentiated from neural crest cells, which were derived from human iPSCs cultured in vitro. These neural crest cells were further grown in supplemented media to promote their differentiation and maturation into mature peripheral sensory neurons (Figure 1A). Single-nucleus RNA sequencing (snRNA-seq) combined with Uniform Manifold Approximation and Projection (UMAP) analysis demonstrated the heterogeneity of the cell population derived from this neuronal differentiation platform, including nociceptors, mechanoreceptors, trigeminal neurons, undefined neurons, astrocytes, oligodendrocytes, and epithelial cells (Figure 1B). Furthermore, the result showed that TRPV1 is highly expressed in nociceptors, mechanoreceptors, trigeminal neurons (Figure 1C).

**Figure 1:** Generation and Characterization of iPSC-Derived Sensory Neurons. **(A)** Schematic diagram and phase contrast images illustrate the generation of sensory neurons from iPSCs via neural crest cell differentiation. Phase contrast mages were captured with a EVOS M100 fluorescence microscope using a 10× objective. Scale bar, 300 μm or 100μm, as indicated. **(B)**. snRNA-seq UMAP analysis results showed the cell population derived from this neuronal differentiation platform included nociceptors, mechanoreceptors, and trigeminal neurons, undefined neurons, astrocytes oligodendrocytes and epithelial cells and the TRPV1 expression in each cell type. **(C)**. snRNA-seq UMAP analysis results showed that TRPV1 is highly expressed in nociceptors, mechanoreceptors, and trigeminal neurons.

The sensory neurons were immunoassayed to examine their expression of peripherin, a marker for peripheral neurons, and ACE2, the receptor for SARS-CoV-2. Our results indicate that peripherin-positive sensory neurons express low but detectable levels of ACE2 (red–upper images) under control conditions. When these sensory neurons were exposed to the Omicron BA.5 virus, originally isolated from COVID-19 patients in Hong Kong (hCoV-19/Hong Kong/HKU-220712-005/2022), at a multiplicity of infection (MOI) of 1.0, ACE2 expression was significantly upregulated in some of the sensory neurons. Additionally, there was noticeable enlargement of the neuronal soma and a marked decrease in peripherin staining in both the soma and processes, indicative of neuronal damage (Figure 2A & 2B). To further investigate the cause of this ACE2 upregulation, neuronal network cultures were exposed to the S1 subunit of the BA.5 spike protein. The cells were extensively washed with PBS, fixed, and ACE2 expression was examined 24 hours post-infection (hpi). The results indicated that the SARS-CoV-2 spike protein binds to sensory neurons, significantly upregulating ACE2 levels (green) within 24 hours of exposure (Figure 2C & 2D).

**Figure 2:** Effect of Exposure to Live Omicron BA.5 Virus or Its Spike Protein S1 Subunit on ACE2 Expression in Peripherin-Positive Sensory Neurons. **(A)** Immunostaining of mature sensory neurons exposed to BA.5 virus for 48 hours. Differentiated neurons were exposed to BA.5.at an MOI of 1 or left mock-exposed (Mock), and after 48 hours The cells were fixed and immunoassayed with a peripherin (green) and ACE2 (red) antibodies and stained with DAPI (Panels A) resulted in increased ACE2 expression, enlarged neuronal soma, and damage to neuronal processes. Images were captured with a confocal microscope using a 20× objective. Scale bar, 50 μm. **(B)** Quantification of ACE2 expression levels in mock- and BA.5 virus-exposed neurons from three images captured from different fields. The levels of red fluorescence per captured image for antibody-stained cells from (A) were quantified using Image J software and normalized against the mock-exposure control x-axis label to red fluorescence. **(C)**. ACE2 expression levels following exposure to the S1 subunit of the BA.5 spike protein. Differentiated neurons were mock-exposed or exposed to S1 (100ng/ml) for 24 or 48 hours, then immunoassayed for S1 (red) and ACE2 (green) and mounted with DAPI (blue) Images were captured with a confocal microscope using a 40× objective. Scale bar, 20 μm. **(D)** Quantification of ACE2 expression in control and S1-exposed neurons. Red ACE2 expression in cells from (C) were assessed as described above (B). Data were analyzed by Student’s t-test in GraphPad Prism 10 and are presented as means ± SDP-values are indicated in the figure: **** p < 0.0001.

### 2. Omicron BA.5 Virus Directly Infects TRPV1-Positive Neurons, Including Olfactory Neurons

To investigate whether specific subtypes of sensory neurons can be directly infected by the BA.5 virus, we generated an BA.5 virus expressing mCherry by inserting the mCherry reporter gene into the F6 fragment of the SARS-CoV-2 BA.5 genome using a circular polymerase extension reaction (CPER) method ^29^.The BA.5 virus expressing mCherry was administered to neuronal cultures at 0.1, 1.0, or 2.0 MOI, and the cultures were incubated at 37°C in 5% CO2. Neurons were observed daily for mCherry fluorescence signals using a fluorescence microscope to identify infected cells. At 24 hpi, mCherry fluorescence was detected in the 1.0 and 2.0 MOI groups (preliminary data not shown). By 48 hpi, mCherry fluorescence was observed in all infection groups, with the 1.0 MOI chosen as the exposure dose for further studies. Our results indicated that approximately 5-10% of neurons became directly infected, with the BA.5 virus predominantly localizing to the neuronal soma and colocalizing with TRPV1. Since TRPV1 is highly expressed in nociceptive, trigeminal neurons (Figures 1C), we aimed to confirm whether TRPV1-positive olfactory neurons are susceptible to SARS-CoV-2 infection. Cell cultures exposed to the BA.5 variant were examined through immunofluorescence analysis using TRPV1-specific and olfactory marker protein (OMP)-specific antibodies. The results demonstrated that olfactory neurons expressed TRPV1 and are susceptible to direct infection by the virus. Additionally, virus exposure led to the translocation of TRPV1 (purple) from the nucleus to the cytosol (Figure 3B). Viral infection was further confirmed through staining with a nucleocapsid protein (NP)-specific antibody, which supported the finding that the BA.5 variant can directly infect sensory neurons (Figure 3C).

**Figure 3:** BA.5 Virus Infection of Sensory Neurons. **(A)** Immunostaining of mCherry-expressing SARS-CoV-2 virus (red) or mock exposed sensory neurons at 48 hours post infection. Peripherin (green) and TRPV1 (purple) staining indicate the total number of Peripherin positive neurons in each field, including those not infected with the mCherry-expressing BA.5 virus. DAPI (blue). Images were captured with a confocal microscope using a 10× objective. Scale bar, 100 μm. **(B)** Immunostaining of mCherry-expressing SARS-CoV-2 virus in neurons together with the olfactory marker protein OMP (green) and TRPV1 (purple) to determine if olfactory sensory neurons can be infected by the BA.5 virus. Scale bars is 40 μm **(C)** Confirmation of viral infection through immunostaining for SARS-CoV-2 BA.5 nucleocapsid protein (NP-green) and peripherin (red) to detect the presence of viral protein within neurons. Images in panels B and C were captured with a confocal microscope using a 40× objective. Scale bars, 40 μm.

### 3. Omicron BA.5 Virus and Its Spike Protein S1 Subunit Induce TRPV1 Translocation from the Nucleus to the Cytosol

TRPV1 activation through various physical and chemical stimuli, including capsaicin, heat, low pH, and toxins, can trigger acute nociceptive pain and neurogenic inflammation. To investigate TRPV1 activation in sensory neurons, we examined the effects of the Omicron BA.5 virus and its S1 spike protein subunit. Sensory neurons infected with BA.5 were analyzed via immunofluorescence with peripherin (green) and TRPV1 (red) specific antibody staining at 24 or 48 hpi. Upon exposure to the virus, the neuronal soma noticeably enlarged, and most peripherin-positive neurons upregulated TRPV1 expression. This upregulation was accompanied by the translocation of TRPV1 from the nucleus to the cytosol. In contrast, mock-exposed neurons predominantly localized TRPV1 in the nucleus, where it colocalized with DAPI (Figure 4A). Figure 4B shows the total fluorescence detected in each cellular compartment, suggesting that exposure to the BA.5 virus resulted in more TRPV1 localized to the cytosol. Treatment with the S1 subunit of the BA.5 virus spike protein (100 ng/mL) in neuronal cultures yielded similar results. Immunostaining with S1 (green) and TRPV1 (red) antibodies showed colocalization of the two proteins, resulting in a yellow signal. The viral S1 subunit induced upregulation of TRPV1 expression and triggered TRPV1 translocation. However, more TRPV1 remained in the nucleus (Figures 4C & D). Furthermore, we compared TRPV1 activation mediated by the S1 spike protein subunits from the Omicron BA.5 and original Wuhan SARS-CoV-2 viruses, with the latter exhibiting stronger pathogenic properties. The Wuhan S1 protein induced a stronger response than the BA.5 S1 protein, leading to increased TRPV1 expression (Figures 4E & F) and enhanced translocation of TRPV1 to the cytosol. This was accompanied by a reduction in TRPV1 retention in the nucleus (Figures 4E & G).

**Figure 4:** Effect of BA.5 Virus and S1 Spike Protein Subunits on TRPV1 Expression. **(A)** Immunostaining of neurons exposed to BA.5 virus or mock-exposed with peripherin (green) and TRPV1 (red) specific antibodies. DAPI (blue) indicates cell nuclei. White arrows indicate the cellular location of TRPV1 in control and virus-exposed cells. Virus exposure induced upregulation of TRPV1 expression, along with TRPV1 translocation from the nucleus to the cytosol. **(B)** Quantification of TRPV1 cellular location and expression levels in response to BA.5 virus. **(C)** Sensory neurons exposed to BA.5 S1 protein, or mock-exposed, stained with antibodies specific for S1 (green) and TRPV1 (red). The arrows showed the increase of TRPV1 trafficking to cytosol in S1-exposed cells, showing a similar effect to TRPV1 translocation induced by in BA.5 virus-exposure. **(D)** Quantification of TRPV1 cellular location and expression levels in response to BA 5 viral S1 proteins. **(E)** Comparison of TRPV1 expression and cellular location in sensory neurons exposed to Omicron BA.5 or Wuhan S1 proteins (immunostaining as for panel A). Images in A-E were captured with a confocal microscope using a 40x (amend objective. Scale bar, 20 μm. **(F)** Quantification of TRPV1 expression levels and cellular location in response to BA.5 virus and Wuhan viral S1 proteins. Data were analyzed by Student’s t-test in GraphPad Prism 10 and are presented as means ±SD. P-values are indicated in the figure: **** p < 0.0001, ** p < 0.01, * p < 0.05.

### 4. Single-Nucleus RNA Sequencing Reveals Sensory Neuron Response to BA.5 Virus Exposure and Neuroprotective Effects of TRPV1 Antagonist Capsazepine

To better understand the impact of SARS-CoV-2 on sensory neurons, we conducted single-nucleus RNA sequencing (snRNA-seq) analysis on sensory neurons infected with the BA.5 virus and compared them to mock-infected controls. The results revealed a significant reduction in the number of sensory neurons in the infected group compared to controls (Figure 5A), whereas glial cells, such as astrocytes, were less affected (Figure 5B). GO enrichment analysis of the sensory neurons showed that the most upregulated molecular functions included active transmembrane transporter activity and virus receptor activity (Figure 5C). Additionally, KEGG enrichment analysis indicated that virus exposure influenced several pathways, including the cAMP signaling pathway, inflammatory mediator regulation of TRP channels, lysosome function, axon guidance, and dopaminergic and glutamatergic synapses (Figure 5D).

**Figure 5:** Sn-RNA Sequencing Analysis of BA.5 Virus Exposure Response: Capsazepine and Mutant BA.5 Virus Alleviate Axon Injury. **(A)**. The ratio of sensory neurons number was lower in BA.5 infected sensory neurons compared with mock infected group **(B)** The ratio of astrocytes remained relatively unchanged between the BA.5 exposure group and the mock-infected group. **(C)**. Sensory neurons GO term enrichment analysis showing the upregulated molecule function between BA.5 exposure and mock control. The numbers within the figure refer to enrichment factor **(D)**. Sensory neurons KEGG enrichment analysis of the impact of KEGG pathways between BA.5 exposure and mock control samples. The numbers within the figure refer to enrichment factor **(E)** Immunostaining of neuronal networks exposed to BA.5 virus, Del-FCS BA.5 virus, or BA.5 co-administered with 10 ng/ml of the TRPV1-antagonist capsazepine with peripherin, TRPV1 and NP antibodies (48 hpi). Merged images show peripherin (green), TRPV1 (red), NP (green) and DAPI (blue) staining. Disappearance of peripherin-positive and processes with NP express straining in images indicates the damage of axons. Del-FCS mutant BA.5 virus exposure had less pronounced impact compared with wild-type BA.5. Images were captured using a confocal microscope using a 20× objective. Scale bar, 100 μm. **(F)** Quantification of peripherin expression levels in neurons exposed to BA.5 with or without capsazepine or Del-FCS mutant. **(G)** Neuronal networks exposed to BA.5 virus with or without 10 ng/ml capsazepine or to Del-FCS mutant BA.5 virus (as above), stained with antibodies specific for TRPV1 (red) or peripherin (green). DAPI (blue) indicates cell nuclei. TRPV1 Inhibitor and Del-FCS mutant virus reduced TRPV1 activation Images were captured with a confocal microscope using a 40× objective. Scale bar, 20 μm. **(H)** Quantification of TRPV1 cytosol: nucleus expression levels in neurons following exposure to BA.5 virus with or without capsazepine treatment or to Del-FCS mutant BA.5 virus, Data were analyzed using Student’s t-test in GraphPad Prism 10 and are presented as means ± SD. P-values are indicated in the figure: **** p < 0.0001, *** p < 0.001, ** p < 0.01, * p < 0.05.

To further investigate the correlation between TRPV1 activation and neuronal degeneration, the BA.5 virus was co-administered with the TRPV1 antagonist capsazepine (10 ng/ml). At 48 hpi, the neuronal networks were immunoassayed with peripherin, viral nucleocapsid protein (NP), and TRPV1 antibodies. The images were analyzed at 20× magnification, focusing on the neuronal processes. The results demonstrated that most sensory neurons lost axons and dendrites, as indicated by peripherin staining, suggesting degeneration following 48 hpi to 1.0 MOI Omicron BA.5 virus. The loss of axonal structures caused by BA.5 virus exposure was attenuated by co-administration of capsazepine. This was further confirmed by viral NP protein staining, which showed that infected neurons retained their processes in the presence of the TRPV1 inhibitor (Figure 5 E&F). Additionally, we assessed TRPV1 activation and axonal damage using the attenuated Del-FCS BA.5.2 mutant virus strain in the sensory neuronal network. This mutant strain, engineered with a deletion at the S1/S2 furin cleavage site (FCS: KSPRRARSVA) within the SYQTQTKSPRRARSVASQSIIA sequence, inhibits viral cleavage and reduces viral entry and replication capacity (unpublished data). The results revealed that, compared to the wild-type virus, the mutant strain had a lesser impact on TRPV1 expression, trafficking, and neuronal process destruction (Figure 5G&H). These observations suggest a relationship between virus entry, replication, pathogenic properties, and TRPV1 activation, with TRPV1 activation promoting axonal damage in neurons. Our results imply that TRPV1 activation could serve as a biomarker for assessing the impact of different SARS-CoV-2 strains on sensory neurons and may represent a potential target for therapeutic intervention.

## Discussion

Infection with SARS-CoV-2 is associated with dysfunction across multiple organ systems. During the acute phase of infection, in addition to respiratory symptoms, some patients experience neurological abnormalities, such as headache, anosmia, and ageusia ^2^. Primary sensory neurons are specialized to detect intense stimuli and serve as the first line of defense against potentially harmful environmental inputs through peripheral sensitization. The susceptibility of sensory neurons, including those in the olfactory system, to SARS-CoV-2 infection underscores the need to explore protective strategies and therapeutic interventions. This is particularly important given that neurological symptoms are among the most common manifestations of long COVID.

Two mechanisms have been proposed to explain the impact of viral infection on olfactory sensory neurons. The direct mechanism suggests that viruses, such as influenza and coronaviruses, can directly infect olfactory receptor neurons, leading to neuronal death and resulting in the loss of the sense of smell ^10^. In contrast, the indirect mechanism proposes that viruses infect cells within the olfactory epithelium, disrupting the epithelial barrier, which in turn affects the underlying olfactory neurons ^8, 9, 30^. The findings of using i*n vitro* experimental system indicate that SARS-CoV-2 can both directly invade olfactory and other sensory neurons and indirectly affect these neurons by promoting TRPV1 activation through exposure to the viral spike protein.

In addition to direct viral infection, we observed that in cultures infected with BA.5, most TRPV1-positive sensory neurons, including olfactory neurons, upregulated TRPV1 expression. Furthermore, these neurons demonstrated translocation of TRPV1 from the nucleus to the cytosol and its insertion into the cell membrane, even in cells that showed no detectable signs of viral infection (Figure 3). TRPV1 is expressed in various cell types, including the central nervous system, nociceptors, trigeminal nerves, and olfactory neurons. Nociceptor neurons are highly specialized to respond to noxious stimuli and relay this information to the central nervous system for appropriate responses. The nose serves as the primary site for viral entry. In mammals, the trigeminal and olfactory systems mediate the detection of volatile chemicals, and these two neural systems interact with one another ^31-33^. Our observations indicate that nociceptors, trigeminal nerves, and olfactory neurons are major targets for SARS-CoV-2 infection among peripheral sensory neurons. Additionally, exposure to SARS-CoV-2 promoted the nuclear-to-cytosolic translocation of TRPV1, leading to axonal destruction.

The damage to sensory neuron axons and dendrites through TRPV1 activation is highly significant, as axonal damage not only causes sensory neuron disorders but also impairs brain activity and cognitive function. Previous studies have shown crosstalk between TRPV1 and the microtubule cytoskeleton at multiple levels ^20^. TRPV1 activation leads to the rapid disassembly of microtubules, which are essential for neuronal migration and neurogenesis. This disruption of dynamic microtubules due to TRPV1 activation profoundly affects axonal growth, morphology, and migration. The underlying molecular mechanisms of SARS-CoV-2-mediated TRPV1 activation still require further investigation. Single-cell transcriptome analysis revealed that viral exposure increases cAMP signaling. The cAMP pathway can modulate TRPV1 activation through Protein Kinase A (PKA), which phosphorylates TRPV1, sensitizing the channel and enhancing its activity. This, in turn, increases transmembrane transporter activities, promoting inflammation and neurodegeneration. We hypothesize that SARS-CoV-2 can function as inflammatory mediator that regulates TRPV1 activation, thereby promoting axonal injury and inflammation. Notably, the use of TRPV1 antagonists conferred a neuroprotective effect. Further delineating the interaction between SARS-CoV-2 and TRPV1 could reveal potential approaches to mitigate sensory neuron loss and other related effects of viral infection. TRPV1 channels are tetrameric membrane proteins, each subunit containing six transmembrane segments (S1-S6), with a pore region located between S5 and S6. The N-terminal region of TRPV1 contains multiple ankyrin repeat domains (ARDs), each comprising 33-residue motifs that form a helix-loop-helix structure. These ARDs facilitate interactions with various intracellular proteins, including regulatory and scaffolding proteins, which can modulate TRPV1 channel gating and trafficking. Although the precise interaction between SARS-CoV-2 and TRPV1 is not yet fully understood, recent research has identified two ankyrin-binding motifs (ARBMs) within the SARS-CoV-2 spike protein. One is in the S1 subunit, which binds to the host cell receptor ACE2, while the other is in the S2 subunit, responsible for mediating the fusion of viral and cellular membranes ^34, 35^. The presence of ARBMs in the SARS-CoV-2 spike protein and ankyrin ARDs in TRPV1 suggests that TRPV1 may play a role in the binding of the SARS-CoV-2 spike protein to cells and potentially act as a host response protein ^36^. We observed that BA.5 S1-mediated TRPV1 translocation from the nucleus to the cytosol was less pronounced than with live virus exposure, with some TRPV1 remaining in the nucleus. We hypothesize that the S2 protein or other viral factors might also be involved in TRPV1 activation. Future research is needed to explore how both the S1 and S2 subunits of the spike protein, along with other viral factors, interact with TRPV1 and the signaling pathways activated by these interactions.

Crosstalk between TRPV1-positive nerve fibers and immune cells is crucial for mediating airway inflammation during respiratory viral infections ^37^. Understanding this interaction could aid in designing more effective, specific therapeutic agents, potentially including analgesics and anti-inflammatory drugs targeting TRPV1. Investigating how TRPV1 activation mediates neuronal inflammation and how this may influence the immune response in SARS-CoV-2 infections will enhance our understanding of the virus’s infectivity and pathogenesis. This study may open new avenues for developing improved diagnostic approaches and therapeutic interventions. Further research into the mechanism of TRPV1-mediated axonal and dendritic degeneration could also help identify strategies to mitigate sensory neuron loss and other related effects of viral infection.

### Experimental Procedures

#### 1. Differentiation of Peripheral Sensory Neurons from iPSCs

The human iPSC line BIONi010-C (Sigma-Aldrich, Cat #66540666) was cultured in mTeSR Plus serum-free stem cell culture medium (Stem Cell Technologies, Cat #85850) at 37°C and 5% CO2. Neural crest cells (NCCs) were generated from iPSCs using Neural Crest Induction Medium (Stem Cell Technologies, Cat #08610) according to the manufacturer’s instructions. NCCs were further subculture at a cell density of 2 × 10^5 cells/cm^2^ in 4-well culture slides and 24-well plates coated in Matrigel gel (Corning, Cat# 354277) for sensory neuron differentiation and maturation using the STEMdiff™ Sensory Neuron Differentiation Kit (Stem Cell Technologies, Cat #100-0341) and the STEMdiff™ Sensory Neuron Maturation Kit (Stem Cell Technologies, Cat #100-0684) respectively Sensory neurons can be maintained in culture for up to 4 weeks.

#### 2. Virus Strain and Propagation

The BA.5 strain (hCoV-19/Hong Kong/HKU-220712-005/2022; GISAID: EPI_ISL_13777658) was isolated from combined nasopharyngeal-throat swabs of COVID-19 patients in Hong Kong. The viral isolate was propagated in VeroE6 cells. When cytopathic effects (CPE) were observed in 80-90% of the cells, the spent culture medium was centrifuged at 1000 × g for 10 minutes at 4°C. The supernatant was collected, aliquoted, and stored at -80°C as a viral stock. The titer of the viral stock was determined using a plaque assay. All experiments involving authentic SARS-CoV-2 adhered to approved standard operating procedures within the Biosafety Level 3 (BSL-3) facility located at the Department of Microbiology, HKU.

#### 3. Recombinant SARS-CoV-2 Generation by Circular Polymerase Extension Reaction (CPER)

Viral RNA was extracted from the BA.5 strain and used as a template for cDNA synthesis using Superscript IV Reverse Transcriptase with random hexamer primers according to the manufacturer’s protocol (Invitrogen, Cat# 18090050). Six fragments covering the SARS-CoV-2 genome (F1-F6) with complementary 30-nucleotide ends were amplified from viral cDNA using high-fidelity PrimeSTAR GXL DNA polymerase (Takara Bio, Cat# R050B) with corresponding pairs of primers (material table). Equimolar amounts (0.1 pmol each) of the resulting six fragments and SARS-CoV-2 linker fragments were assembled into a circular full-length cDNA using PrimeStar GXL DNA polymerase under the following cycling conditions: initial denaturation at 98°C for 1 minute, followed by 35 cycles of denaturation at 98°C for 10 seconds, annealing at 60°C for 10 seconds, and extension at 68°C for 15 minutes, followed by a final extension at 68°C for 15 minutes.

#### 4. Generation of mCherry-expressing BA.5 Virus

The mCherry gene was inserted into BA.5 fragment F6 and assembled together with BA.5fragments F1-F5 and linker fragments using CPER. The resulting CPER product was transfected into HEK293T cells using Lipofectamine LTX with Plus Reagent (Invitrogen, Cat# 15338100), following the manufacturer’s protocol. Six hours post-transfection, the cells were trypsinized and transferred to a confluent monolayer of Vero E6 cells in a T25 flask for virus propagation to generate the viral stock.

#### 5. Generation of Del-FCS BA.5 Virus

To generate Del-FCS BA.5 virus, the sequence for the furin cleavage site (FCS) located in BA.5 fragment F5 was deleted through CPER technique. The F5 fragment with the deleted FCS was assembled with BA.5 F1-F4 and F6 fragments using CPER. The resulting CPER product was transfected into HEK293T cells using Lipofectamine LTX with Plus Reagent, following the manufacturer’s protocol. At six hours post-transfection, the cells were trypsinized and transferred to a confluent monolayer of Vero E6 cells in a T25 flask for virus propagation to generate the viral stock.

#### 6. Exposure of Sensory Neuronal Networks to Virus and its S1 protein

BA.5 and Wuhan spike protein S1 subunit (abcam, Cat# ab273068, and Acrobiosystems, Cat# S1N-C52Hy) were diluted in Sensory Neuron Maturation Medium to 100ng/mL. Neural crest cells, seeded at a density of 2 × 10^5 cells/cm^2^, were cultured in 4-well culture slides and 24-well plates coated with Matrigel. Following neuronal differentiation and maturation, the peripheral sensory neurons were exposed to SARS-CoV-2 BA.5, mCherry-expressing BA.5, or Del-FCS BA.5 virus diluted in Sensory Neuron Maturation Medium (from the STEMdiff™ Sensory Neuron Maturation Kit) at MOIs of 0.1, 1, or 2. An MOI of 1 corresponds to a 1:1 ratio of cells to virus particles. Mock-exposed cultured neurons served as controls. Capsazepine (MedChemExpress, Cat #HY-15640) at 10 ng/ml was included in some BA.5-exposed cultures for TRPV1 inhibition studies. Cultures were incubated at 37°C with 5% CO2 for 24 or 48 hours. After incubation, the supernatant was removed, and the cells were washed with PBS prior to being processed for immunofluorescence staining. Indicate here the number of replicates for each condition.

#### 7. Immunofluorescence Staining and Microscopy

Neurons were fixed with 4% paraformaldehyde in PBS for 15 minutes, washed with PBS, and permeabilized with 0.1% Triton-X in PBS for 10 minutes. The neurons were blocked in 1% BSA for 1 hour at room temperature and incubated with primary antibodies against peripherin (Invitrogen, Cat# MA3-16724, and abcam, Cat# ab246502), TRPV1 (Abcam, Cat #ab305299), two ACE2 (Invitrogen, Cat #MA5-32307, and abcam, Cat# ab89111), OMP (Santa Cruz Biotechnology, Cat #sc390543,SARS-CoV-2 S1 subunit (abcam, Cat# ab283942), SARS-CoV-2 nucleocapsid protein (Sino Biological, Cat# 40143-MM05) separately or in various combinations, overnight at 4°C. After primary antibody incubation, the cells were washed and incubated with secondary antibodies for 1 hour at room temperature. Coverslips were mounted with VECTASHIELD PLUS Antifade Mounting Medium with DAPI (Vector Laboratories, Cat #H-1200-10).

Phase contrast images were captured using EVOS M5000 Imaging System (Thermo Fisher Scientific, USA). Immunostained images were captured using a Zeiss LSM 880 confocal microscope. Representative images for each experimental condition were chosen for inclusion in figures. For expression analysis data were collected from three independent images and analyzed using ImageJ software (https://imagej.nih.gov/ij/). For ACE2 quantification, the cell body is cropped and measured for the mean fluorescence in the cropped area. The average mean fluorescence of the mock-exposed cells was used as a benchmark to calculate the fold change of mean fluorescence of BA.5-infected cells or S1 glycoprotein-exposed cells. For TRPV1 quantification, the nucleus and cytosol of selected cells were cropped to quantify their integrated density separately to showcase the distribution and expression of TRPV1 within the cell. The ratio of nuclear mean fluorescence to cytosolic mean fluorescence was also calculated to demonstrate the activation of TRPV1 in different treatment groups. For peripherin quantification, 5 fields from cells clusters (each field including cell body and processes) were selected for each group, and cropped to measure the integrated density of peripherin expression. Measured fluorescence data was subjected to one-way ANOVA and Student’s t-test; a p-value < 0.05 was considered significant. Correlations were determined by calculating Pearson coefficients. All statistical analyses were conducted using GraphPad Prism 10 (GraphPad Software, Inc.).

#### 8 Single-nuclei RNA Sequencing (snRNA-seq) Sample Preparation

Two sensory neuron cultures were exposed to 1.0 MOI of the BA.5 virus for 24 hours, while two mock-exposed cultures served as controls, resulting in four samples (#BA.5.1, #BA.5.2, #Mock1, and #Mock2). Neurons were incubated in Accutase (Stem Cell Technologies, Cat #07920) at 37°C for 7 minutes. Detached cells were centrifuged at 300 xg for 5 minutes, and the resulting cell pellet was resuspended in 200 µL of pre-chilled Lysis Buffer (10 mM Tris-HCl, pH 7.4, 10 mM NaCl, 3 mM MgCl, 0.025% Nonidet P40 Substitute in PBS), followed by a 1-minute incubation on ice. Next, 800 µL of pre-chilled Nuclei Wash Buffer (1% BSA, 0.2 U/µL RNase Inhibitor in PBS) was added to the lysate, and the mixture was centrifuged at 500 xg for 10 minutes. The resulting nuclei pellet was washed with 1 mL of Nuclei Wash Buffer, centrifuged again, and resuspended in 1 mL of the same buffer. The nuclei were then immediately fixed using the Chromium Next GEM Single Cell Fixed RNA Sample Preparation Kit (10X Genomics, Cat #1000414), following the manufacturer’s instructions. Fixed nuclei samples were intermittently evaluated for count and quality using the LUNA dual fluorescence cell counter (Logos Biosystems).

The fixed nuclei samples were sequenced using the Single Cell Gene Expression Flex for Multiplexed Samples (10X Genomics) protocol. Following Probe Hybridization, uniquely barcoded nuclei were counted and pooled in equal numbers, adhering to the guidelines in the 10X Genomics Chromium Fixed RNA Profiling Multiplexed Samples Pooling Workbook Rev B (CG000565) ^38^. The pooled nuclei were washed, targeting 20,000 cells per sample for partitioning into GEMs on Chip Q, GEM barcoding, GEM recovery, pre-amplification, and library construction. Sequencing libraries were prepared in accordance with the User Guide (CG000527) ^39^. Libraries were sequenced on an Illumina NovaSeqX platform using paired-end dual-indexing with a read depth of approximately 300 million reads per sample and a 2×150 read length. Demultiplexing of sequencing libraries was performed with bcl2fastq (Illumina), and FASTQ files were processed using Cell Ranger v7.1.0 (10X Genomics) with the GRCh38-2020-A reference genome.

### snRNA-seq Data Analysis

#### Quality control of scRNA-seq data

Raw reads were aligned to the human genome (hg38), and cells were identified using the *Cell Ranger count* pipeline (v7.1.0). Cells with fewer than 200 detected genes, more than 8,000 detected genes, or a mitochondrial contamination rate exceeding 10% were filtered out as low-quality cells. To further reduce ambient RNA contamination from droplets, the *SoupX* R package (v1.6.1) was applied with default settings. Additionally, potential doublets were removed using the *DoubletFinder* R package (v2.0.3) with default parameters. Quality control and normalization of the scRNA-seq data were subsequently performed using the *Seurat* R package (v4.3.0).

#### Cell clustering and cell-type annotation

The gene expression matrices from the filtered cells were normalized to the total UMI counts per cell and transformed to the natural log scale. Highly variable genes (HVGs) were identified using the *FindVariableFeatures* function, selecting the top 2000 HVGs from the corrected expression matrix. The data were then centered and scaled after regressing out cell cycle effects (S and G2M scores were calculated using the *CellCycleScoring* function in *Seurat*). Principal component analysis (PCA) was performed on the HVGs using the *RunPCA* function. To correct for batch effects, the *RunHarmony* function was applied with default parameters.

Dimensionality reduction and k-nearest neighbor graph construction (k=20) were conducted using the *FindNeighbors* function, based on Euclidean distances in the 50-dimensional PC space. Cell clusters were identified using the Louvain-Jaccard graph-based method, with the clustering resolution set to 0.2 via the *FindClusters* function. For visualization, the *RunUMAP* function was employed to reduce high-dimensional data into a two-dimensional (2D) representation using dimensions 1–20.

Cluster-specific marker genes were identified using the *FindAllMarkers* function with default parameters, and significance in gene expression differences was determined by the Wilcoxon rank-sum test with Bonferroni correction. Cell types were manually annotated based on cluster markers. To determine sample composition by cell type, the number of cells for each type in each sample was calculated, normalized to the total number of cells per sample, and scaled to 100% for each cell type.

#### Cell-type subclustering

For the major cell types, such as astrocytes and neurons, we repeated the previously described steps (normalization, dimensionality reduction, and clustering) to identify subclusters. These subclusters were subsequently annotated as distinct specific cell subtypes. To determine the genes uniquely expressed in each subcluster, we utilized the *FindAllMarkers* function in *Seurat* with default parameters. To calculate the composition of different groups within each subcluster, the number of cells from each group within each subcluster was counted and analyzed.

#### GO and KEGG enrichment analysis

Enrichment scores (p-values) for selected GO/KEGG annotations were calculated using the *clusterProfiler* R package (v4.2.2) through a hypergeometric statistical test with a significance threshold of 0.05. The Benjamini-Hochberg method was applied to estimate the false discovery rate (FDR). Enrichment analysis was performed on the input gene lists, with all genes listed in the *org*.*Hs*.*eg*.*db* database serving as the background. Finally, the dot*plot* function was used for visualization of the results.

## Supporting information

manuscript

## Author Contributions

**Vanessa Co**: Performed most of the experiments except for those involving the use of live virus exposure, curated the data, prepared the manuscript. **Siwen Liu**: Generated mutant virus, conducted live virus exposure experiments. **Rachel Chun-Yee Tam**: conducted live virus exposure experiments. **Bobo Wing-Yee Mok**: conducted live virus exposure experiments. **Honglin Chen**: Provided funding and resource support, conceptualized the study, edited the manuscript. **Yiling Hong**: Conceptualized the study, designed methodology, supervised the research, prepared the manuscript.

## Limitations of our study

The limitations of our study are that we haven’t identified the specific pathways involved in TRPV1 translocation and its role in axonal degeneration. On going single cell analysis with a TRPV1 knockout cell line would provide a better understanding of the mechanisms regulating TRPV1 activation and its role in virus infection. Furthermore, we plan to conduct an *in vivo* study to further examine the role of TRPV1 in inflammation response.

## Resource Availability

Single cell RNA-seq data had been deposited to NCBI bank and will be released upon the acceptance of this manuscript. Any additional information required to reanalyze the data reported in this paper is available from the lead contact upon request.

## Acknowledgements

This work is supported by the InnoHK-Health program, Innovation Technology Commission, Hong Kong SAR, PR China. The authors would like to thank Libby Liao and Dr. Ma Sidi from Accuramed Technology (Guangzhou, China) Limited for their invaluable assistance with snRNA-seq bioinformatics analysis.

## Notes

### Competing Interest Statement

The authors have declared no competing interest.

