## Supplementary material for "SARS-CoV-2 Infects Peripheral Sensory Neurons and Promotes Axonal Degeneration via TRPV1 Activation": manuscript

#### Graphical abstract

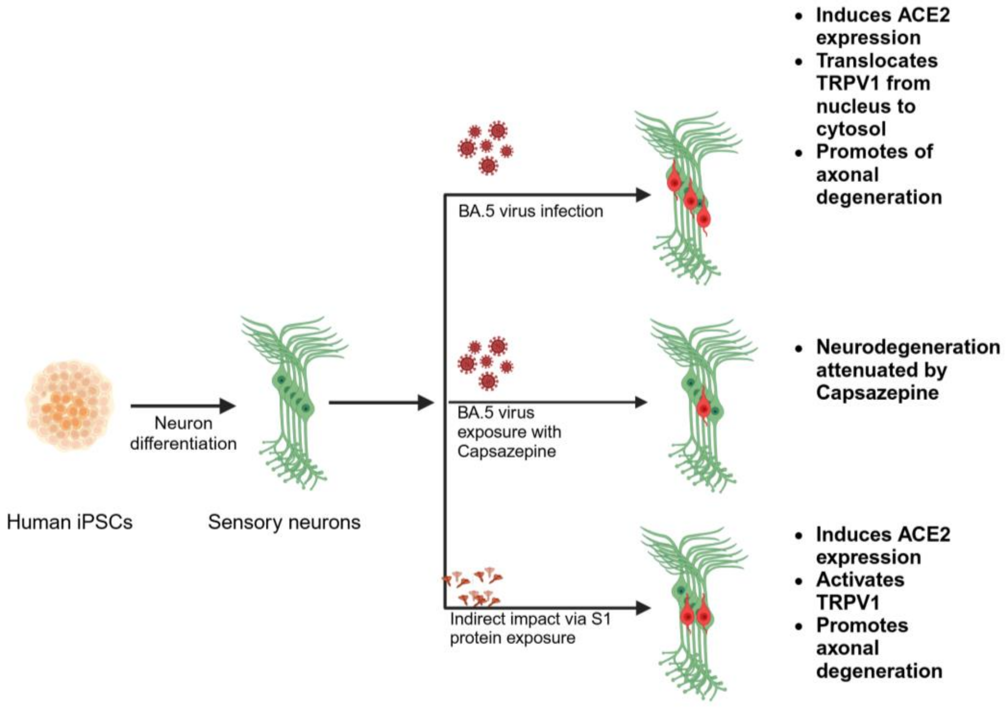


### SARS-CoV-2 Infects Peripheral Sensory Neurons and Promotes Axonal Degeneration via TRPV1 Activation

**Figure 1**

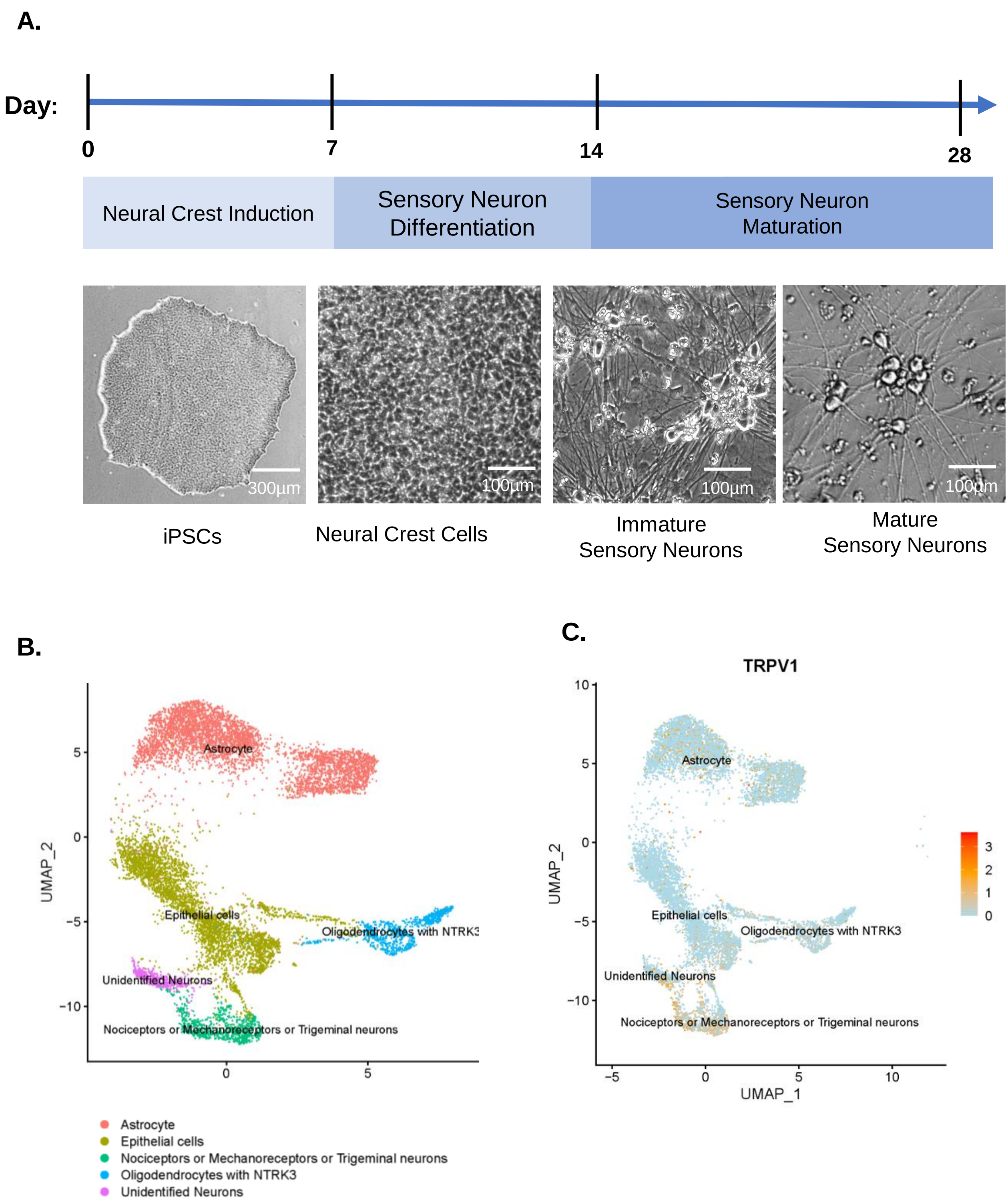
A.
Day:
0
7
14
28
Neural Crest Induction
Sensory Neuron Differentiation
Sensory Neuron Maturation
300µm
100µm
100µm
100µm
iPSCs
Neural Crest Cells
Immature Sensory Neurons
Mature Sensory Neurons
B.
UMAP_2
Astrocyte
Epithelial cells
Oligodendrocytes with NTRK3
Unidentified Neurons
Nociceptors or Mechanoreceptors or Trigeminal neurons
Astrocyte
Epithelial cells
Nociceptors or Mechanoreceptors or Trigeminal neurons
Oligodendrocytes with NTRK3
Unidentified Neurons
C.
TRPV1
UMAP_2
UMAP_1
Astrocyte
Epithelial cells
Oligodendrocytes with NTRK3
Unidentified Neurons
Nociceptors or Mechanoreceptors or Trigeminal neurons

Figure 2

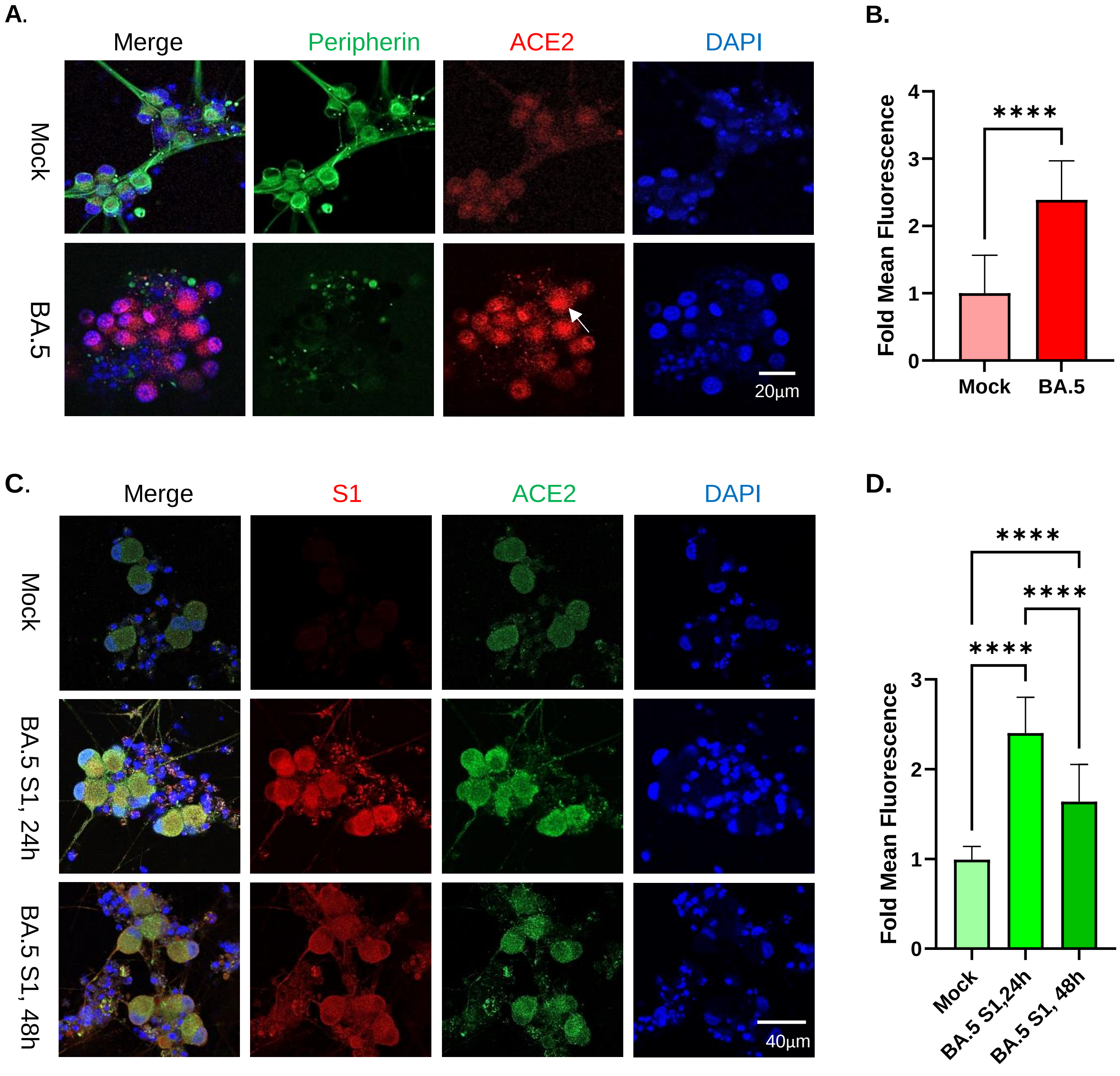

A.
Merge
Peripherin
ACE2
DAPI
Mock
BA.5
20µm
B.
Fold Mean Fluorescence
0
1
2
3
4
****
Mock
BA.5
C.
Merge
S1
ACE2
DAPI
Mock
BA.5 S1, 24h
BA.5 S1, 48h
40µm
D.
Fold Mean Fluorescence
0
1
2
3
****
****
****
Mock
BA.5 S1,24h
BA.5 S1, 48h

**Figure 3.**

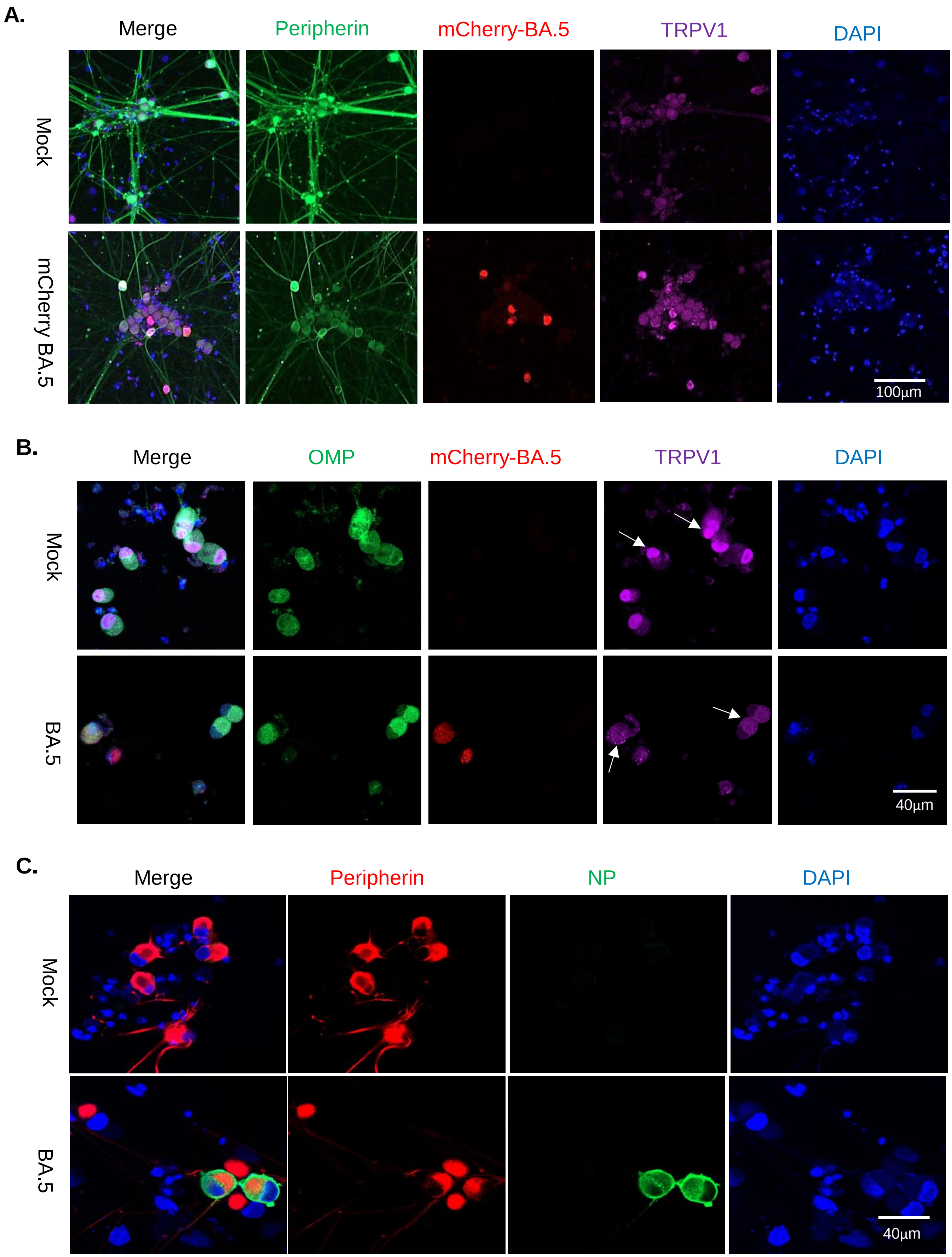
A.
Merge
Peripherin
mCherry-BA.5
TRPV1
DAPI
Mock
mCherry BA.5
100µm
B.
Merge
OMP
mCherry-BA.5
TRPV1
DAPI
Mock
BA.5
40µm
C.
Merge
Peripherin
NP
DAPI
Mock
BA.5
40µm

**Figure 4**

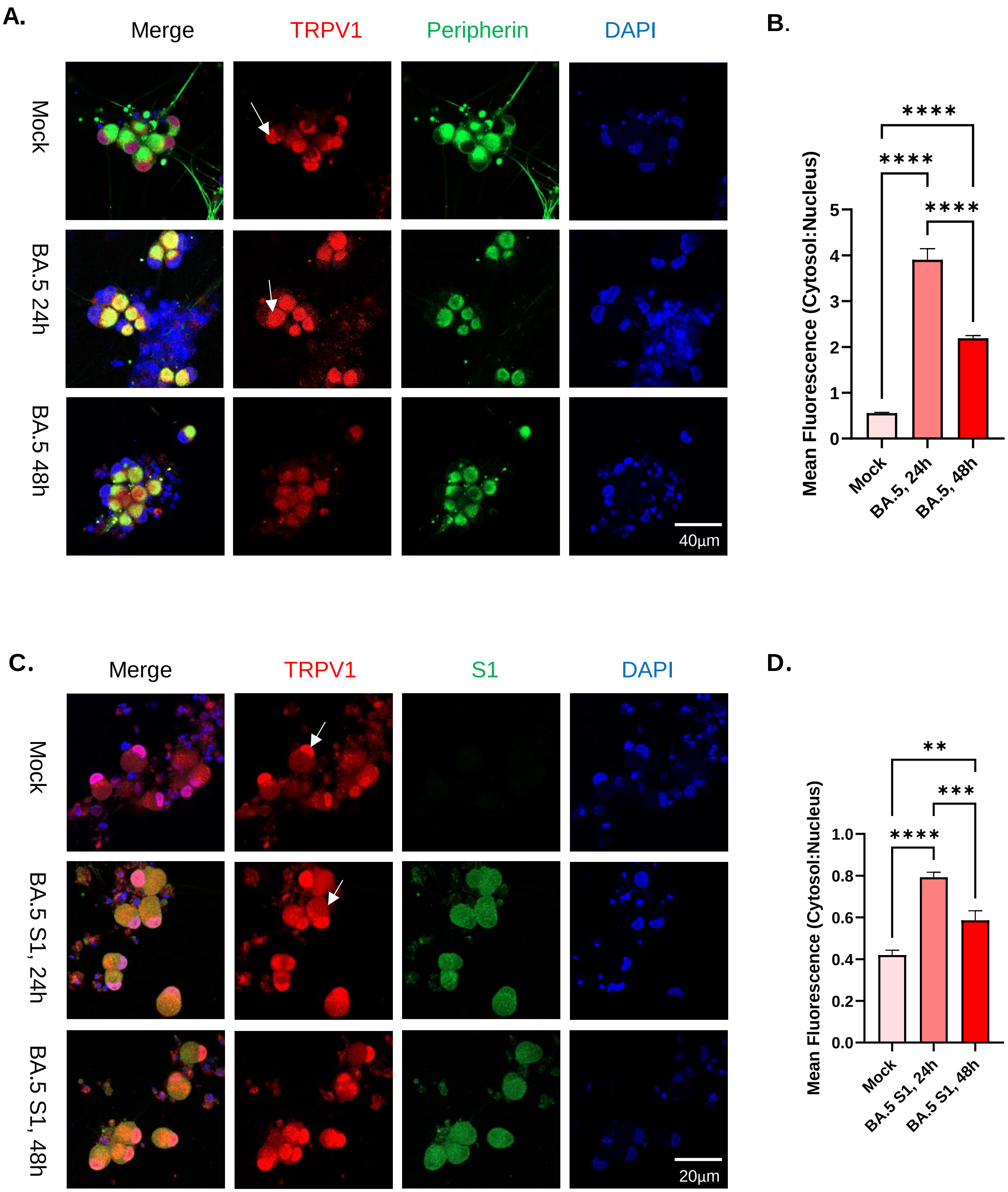

A.
Merge
TRPV1
Peripherin
DAPI
Mock
BA.5 24h
BA.5 48h
40µm
B.
Mean Fluorescence (Cytosol:Nucleus)
0
1
2
3
4
5
Mock
BA.5, 24h
BA.5, 48h
****
****
****
C.
Merge
TRPV1
S1
DAPI
Mock
BA.5 S1, 24h
BA.5 S1, 48h
20µm
D.
Mean Fluorescence (Cytosol:Nucleus)
0.0
0.2
0.4
0.6
0.8
1.0
Mock
BA.5 S1, 24h
BA.5 S1, 48h
**
***
****

E.

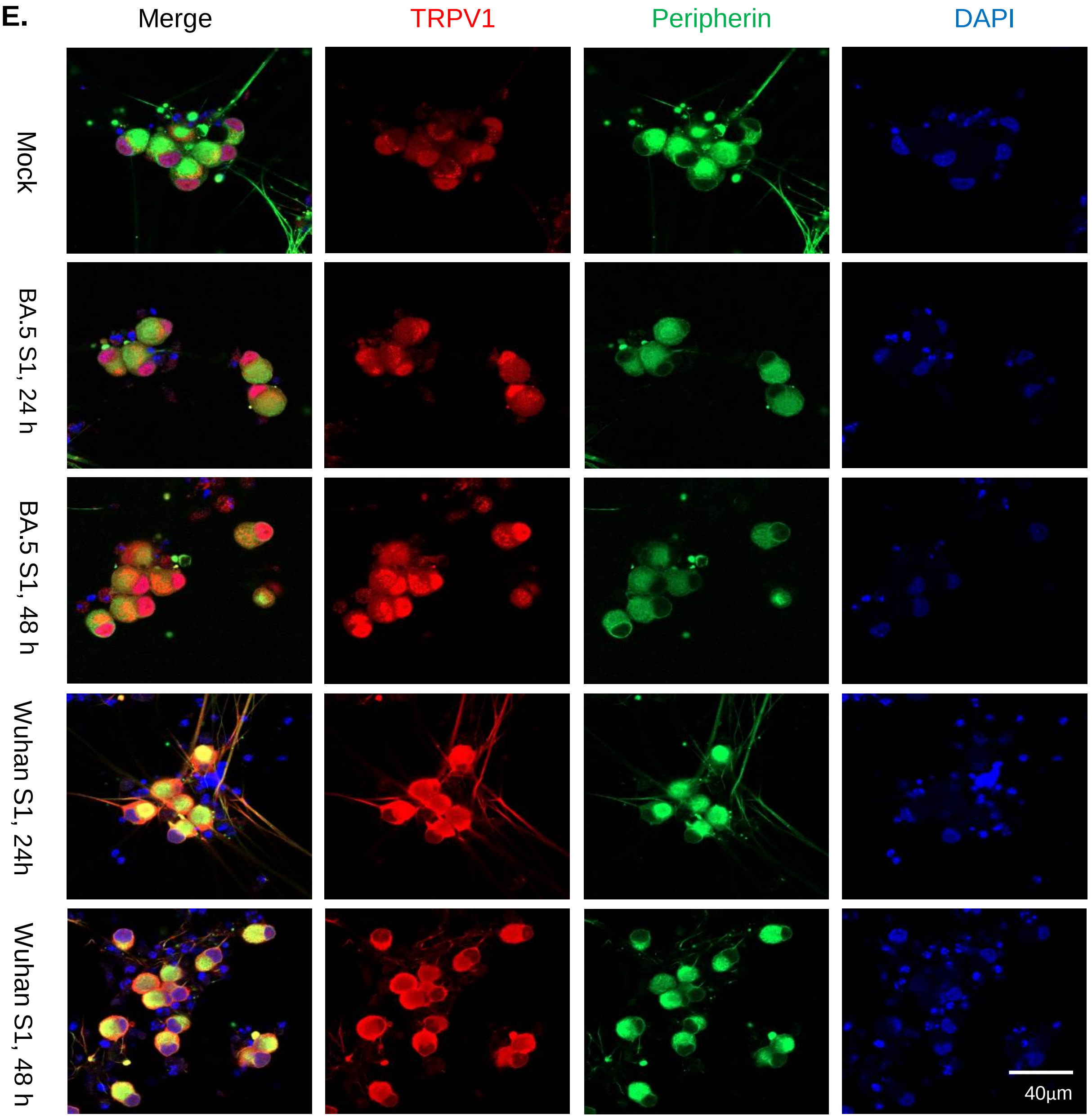

Merge
TRPV1
Peripherin
DAPI
Mock
BA.5 S1, 24 h
BA.5 S1, 48 h
Wuhan S1, 24h
Wuhan S1, 48 h
40μm


F.

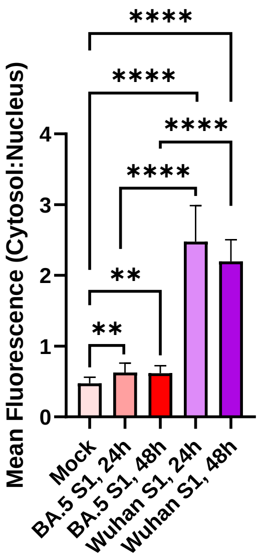

Mean Fluorescence (Cytosol:Nucleus)
0
1
2
3
4
Mock
BA.5 S1, 24h
BA.5 S1, 48h
Wuhan S1, 24h
Wuhan S1, 48h
****
****
****
****
**
**

**Figure 5.**

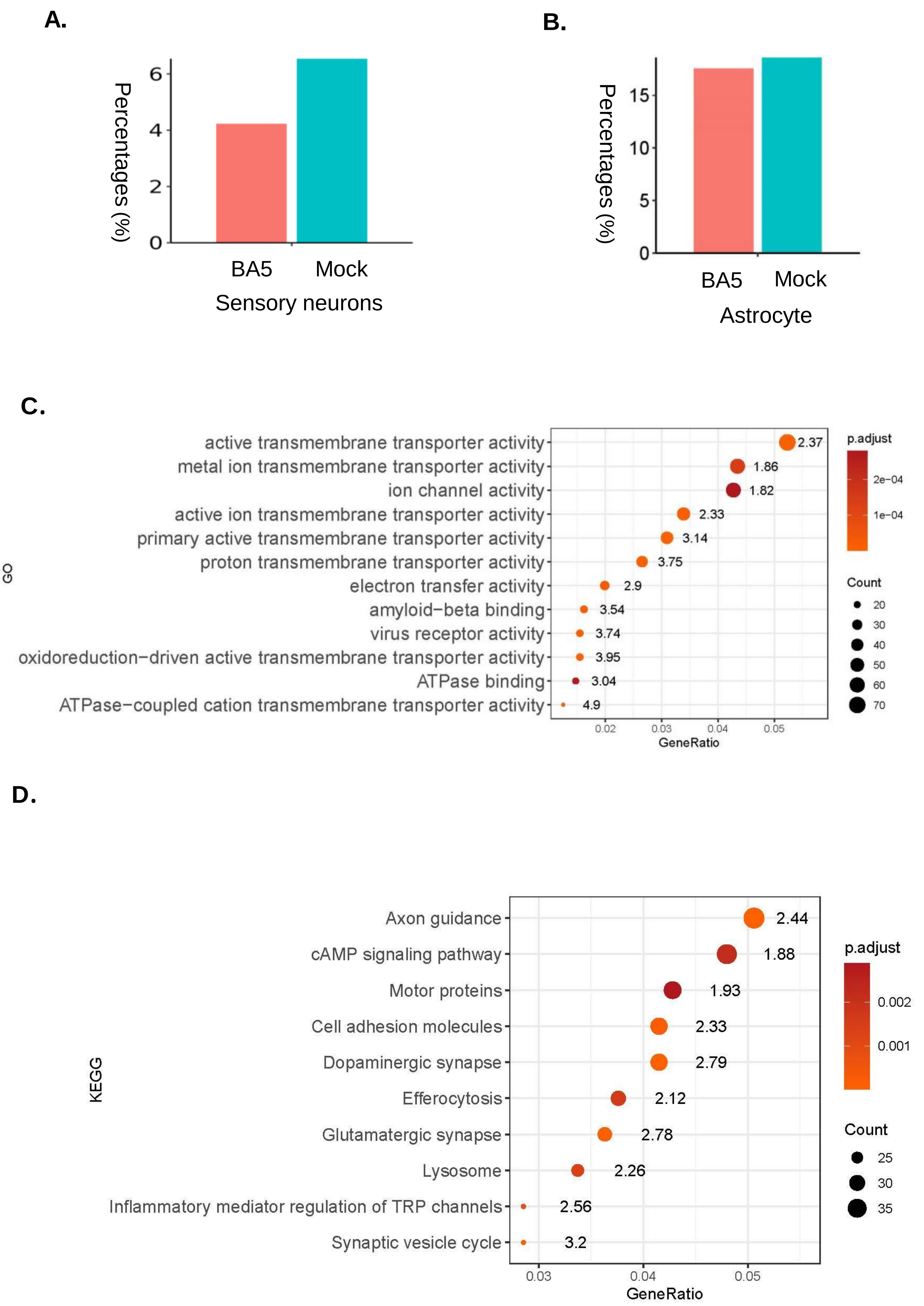
A.
B.
Percentages (%)
0
2
4
6
BA5
Mock
Sensory neurons
0
5
10
15
Astrocyte
C.
GO
active transmembrane transporter activity
metal ion transmembrane transporter activity
ion channel activity
active ion transmembrane transporter activity
primary active transmembrane transporter activity
proton transmembrane transporter activity
electron transfer activity
amyloid−beta binding
virus receptor activity
oxidoreduction−driven active transmembrane transporter activity
ATPase binding
ATPase−coupled cation transmembrane transporter activity
2.37
1.86
1.82
2.33
3.14
3.75
2.9
3.54
3.74
3.95
3.04
4.9
0.02
0.03
0.04
0.05
GeneRatio
p.adjust
2e−04
1e−04
Count
20
30
40
50
60
70
D.
KEGG
Axon guidance
cAMP signaling pathway
Motor proteins
Cell adhesion molecules
Dopaminergic synapse
Efferocytosis
Glutamatergic synapse
Lysosome
Inflammatory mediator regulation of TRP channels
Synaptic vesicle cycle
2.44
1.88
1.93
2.33
2.79
2.12
2.78
2.26
2.56
3.2
0.03
0.04
0.05
GeneRatio
p.adjust
0.002
0.001
Count
25
30
35

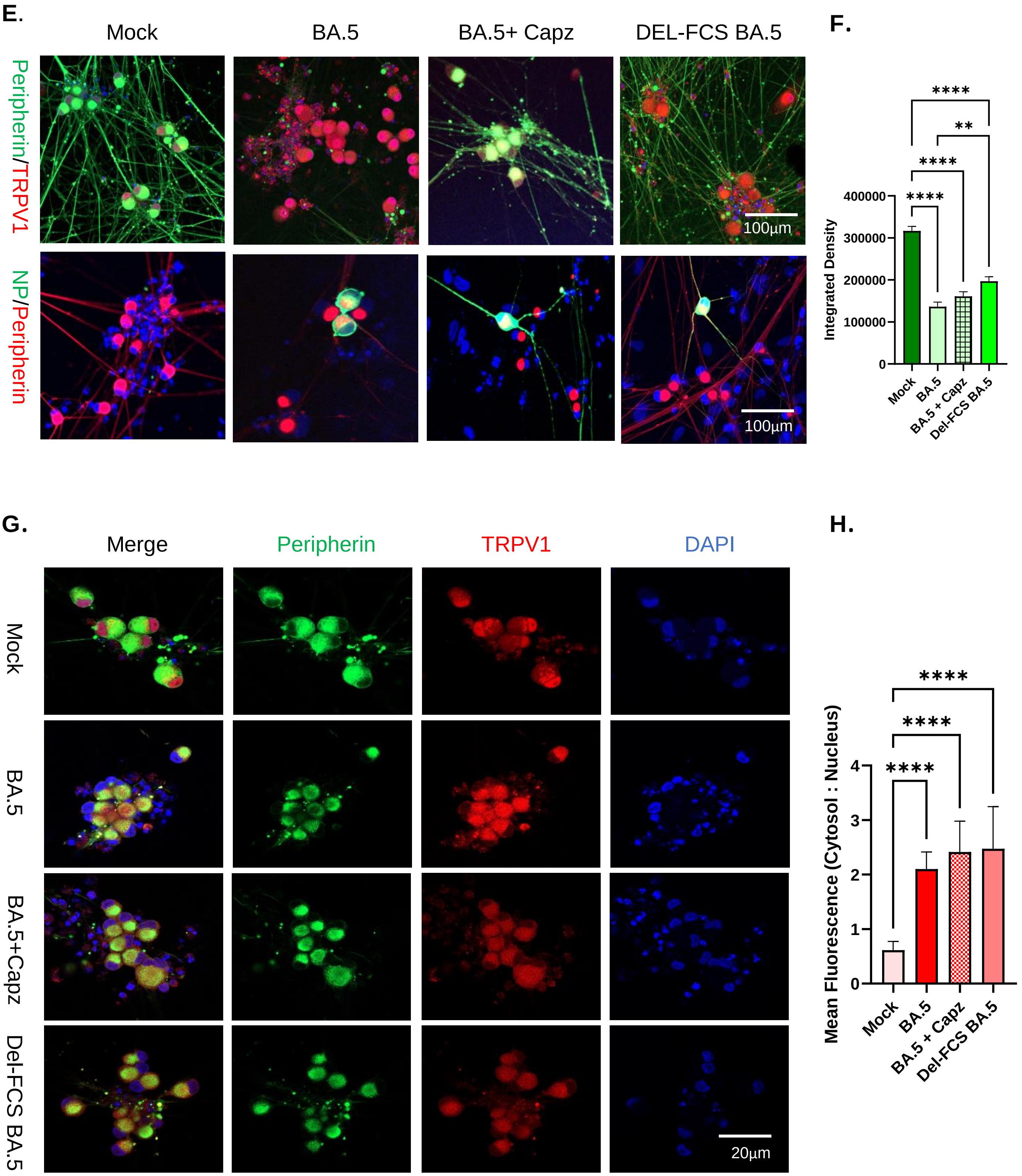

E.
Mock
BA.5
BA.5+ Capz
DEL-FCS BA.5
Peripherin/TRPV1
NP/Peripherin
100µm
100µm
F.
Integrated Density
400000
300000
200000
100000
0
****
**
****
****
Mock
BA.5
BA.5 + Capz
Del-FCS BA.5
G.
Merge
Peripherin
TRPV1
DAPI
Mock
BA.5
BA.5+Capz
Del-FCS BA.5
20µm
H.
Mean Fluorescence (Cytosol : Nucleus)
4
3
2
1
0
****
****
****
Mock
BA.5
BA.5 + Capz
Del-FCS BA.5
